# Gut commensal *Enterocloster* species host inoviruses that are secreted *in vitro* and *in vivo*

**DOI:** 10.1101/2022.10.09.511496

**Authors:** Juan C. Burckhardt, Derrick H.Y. Chong, Nicola Pett, Carolina Tropini

**Affiliations:** Department of Microbiology and Immunology, University of British Columbia, Vancouver, Canada; School of Biomedical Engineering, University of British Columbia, Vancouver, Canada; Humans and the Microbiome Program, Canadian Institute for Advanced Research (CIFAR), Toronto, Canada

**Keywords:** Bacteriophages, Filamentous phages, Inoviruses, Human gut phageome, Human microbiome, Phage-host interactions

## Abstract

**Background:** Bacteriophages in the family *Inoviridae*, or inoviruses, are under-characterized phages previously implicated in bacterial pathogenesis by contributing to biofilm formation, immune evasion, and toxin secretion. Unlike most bacteriophages, inoviruses do not lyse their host cells to release new progeny virions; rather, they encode a secretion system that actively pumps them out of the bacterial cell. To date, no inovirus associated with the human gut microbiome has been isolated or characterized.

**Results:** In this study, we utilized *in silico, in vitro* and *in vivo* methods to detect inoviruses in bacterial members of the gut microbiota. By screening a representative genome library of gut commensals, we detected inovirus prophages in *Enterocloster spp*. and we confirmed the secretion of inovirus particles in *in vitro* cultures of these organisms using imaging and qPCR. To assess how the gut abiotic environment, bacterial physiology, and inovirus secretion may be linked, we deployed a tripartite *in vitro* assay that progressively evaluated growth dynamics of the bacteria, biofilm formation, and inovirus secretion in the presence of changing osmotic environments. Counter to other inovirus-producing bacteria, inovirus production was not correlated with biofilm formation in *Enterocloster spp*. Instead, the *Enterocloster* strains’ inoviruses had heterogeneous responses to changing osmolality levels relevant to gut physiology. Notably, increasing osmolality induced inovirus secretion in a strain-dependent manner. We confirmed inovirus secretion in a gnotobiotic mouse model inoculated with individual *Enterocloster* strains *in vivo* in unperturbed conditions. Furthermore, consistent with our *in vitro* observations, inovirus secretion was regulated by a changed osmotic environment in the gut due to osmotic laxatives.

**Conclusion:** In this study, we report on the detection and characterization of novel inoviruses from gut commensals in the *Enterocloster* genus. Together, our results demonstrate that human gut-associated bacteria can secrete inoviruses and begin to elucidate the environmental niche filled by inoviruses in commensal bacteria.

## Background

Bacteriophages, or phages, are viruses that infect prokaryotic organisms, and they are a major driver of bacterial dynamics in gut-associated microbial ecosystems [1]. Phages from the Inoviridae family (Caudovirales order), known as inoviruses, are widespread throughout most microbial habitats, including human-associated microbial communities [2,3]. Inoviruses have unique filamentous morphologies, circular single-stranded DNA genomes of 4-12kb, and a distinctive lysogenic life cycle [2,4–6]. Unlike other chromosomally integrated bacteriophages that eventually cause host lysis to release infectious progeny, inoviruses encode a secretion system that actively pumps progeny out of the infected bacterium, leading to a chronic and non-lethal infection of the bacterial host [4]. Only 54 inoviruses have been characterized in Gram-negative organisms [7], and notably in pathogens such as *Vibrio cholerae* and *Pseudomonas aeruginosa* [8]. Conversely, in Gram-positive bacteria, only two inovirus hosts have been reported—*Propionibacterium freudenreichii* and *Clostridium beijerincki* [9–11], which are mainly found in dairy and soil, respectively [12,13].

Like other lysogenic phages, inoviruses in their prophage (chromosomally integrated) form can confer unique benefits to their bacterial host, offsetting the burden they may impose with their chronic infection [5]. For example, genes carried within inovirus genomes can affect aspects of biological systems, such as host motility, growth dynamics, biofilm formation and virulence [7,8]. In addition, because of their unique lifestyle that avoids host cell death, inoviruses are effective vectors that can laterally transmit genes to other bacteria [5]. A well-studied case of this is the inovirus CTXΦ, which contains the cholera toxins in its genome and can be laterally transmitted from virulent to non-virulent *V. cholerae* [14]. Infection by CTXΦ leads to a toxin-producing phenotype [14], which induces severe diarrhea in humans, resulting in the transmission, survival, and reproduction of both phage and bacteria [15]. Interestingly, beyond prophage-encoded genes, inovirus particles can also confer benefits to the bacterial host. For example, in Pf-4, the inovirus that infects *Pseudomonas aeruginosa*, secreted virions are implicated in biofilm formation by promoting matrix crystallization [16] and facilitate bacterial infection by activating the immune system and exhausting its responses at the onset of mounting an infection [17].

Despite recent advances in understanding inoviruses in the context of pathogenesis, the implications of inovirus presence in the microbiome have not been well studied. While inoviruses have been detected in the gut, thus far they have not been isolated. However, with the recent developments of virome research and of bioinformatic tools to predict inoviruses, the exploration of this phage family has become more accessible. Inoviruses can affect their bacterial hosts phenotypically, are important vectors for horizontal gene transfer, and can have immunogenic properties. Therefore, dissecting the roles that inoviruses play in the gut could reveal important new biological insights for host-associated bacterial communities and thus for the health and disease of the host organism.

Here, we show a novel characterization of inoviruses from a library of human gut commensal bacteria. We first screened a library of gut bacteria spanning 54 genera and 33 families, which revealed putative inoviruses in members of the genera *Enterocloster* and *Hungatella* — adding 5 new species to only two other reported Gram-positive bacteria capable of secreting inoviruses. We then characterized the secretion of *Enterocloster spp*. inovirus particles under different osmolalities *in vitro* using molecular and imaging methods. We observed species- and strain-dependent responses to perturbations which affected both inovirus secretion and host biofilm formation. Lastly, to investigate inovirus secretion in a more physiologically relevant context, we quantified inovirus production in a mouse model of osmotic diarrhea, confirming that *in vivo* inovirus secretion patterns were consistent with those measured *in vitro*. Our study highlights a previously unknown niche that inoviruses fill in the gut microbiota and opens doors to new studies elucidating their role in shaping complex microbial dynamics in the gut.

## Results

### Uncharacterized inovirus candidates are predicted in commensal gut-associated bacteria

While inoviruses have primarily been studied in the context of pathogens [8,15,18–20], they are globally prevalent phages that have been found across both environmental and host-associated ecosystems through genomics and metagenomics approaches [2,3,21,22]. We sought to investigate which host-associated bacterial species are infected by inoviruses using a genomic approach. To predict inovirus genomes in gut bacteria, we used a previously published bioinformatics pipeline called Inovirus Detector [2]. Briefly, the Inovirus Detector pipeline uses a hidden Markov model similarity search to detect the inovirus *pI* gene, an ATPase part of the extrusion machinery encoded by the phage genome. Importantly, this is the only gene highly conserved among published inovirus genome sequences [2]. Then, the program implements a random forest classifier to compare the surrounding genomic regions flanking the *pI* gene to a curated inovirus database containing distinctive inovirus features. We used this pipeline to screen the genomes of 163 bacteria known to be prevalent in the gut microbiota, representing 130 species and spanning 6 phyla (Figure S1 and Table S1) [23].

Of the 54 bacterial genera screened with Inovirus Detector, we found predicted inovirus genomes in the *Enterocloster* and *Hungatella* genera (previously known as *Clostridium). Enterocloster* and *Hungatella spp*. are anaerobic Gram-positive spore formers that are well-represented in the gut microbiome [24]. 6 out of 7 *Enterocloster* strains and 1 out of 2 *Hungatella* strains in our library were identified as containing the *pI* gene (Figure 1A): *Enterocloster bolteae, Enterocloster clostridioformis, Enterocloster citroniae, Enterocloster aldenensis* and *Hungatella hathewayi* (Table S2) [24]. The inoviruses we predicted had a genome size ranging between 6.5-8.5 kbp, except for *E. aldenensis*, which contained a 16 kbp inovirus genome. Each inovirus genome included a conserved *pI* homolog in addition to 10-16 other open reading frames (ORF). Furthermore, all inovirus genomes were flanked with direct repeats, in line with typical attachment sites where phage genomes integrate through homologous or site-specific recombination into bacterial genomes [25–27].

**Figure 1.**
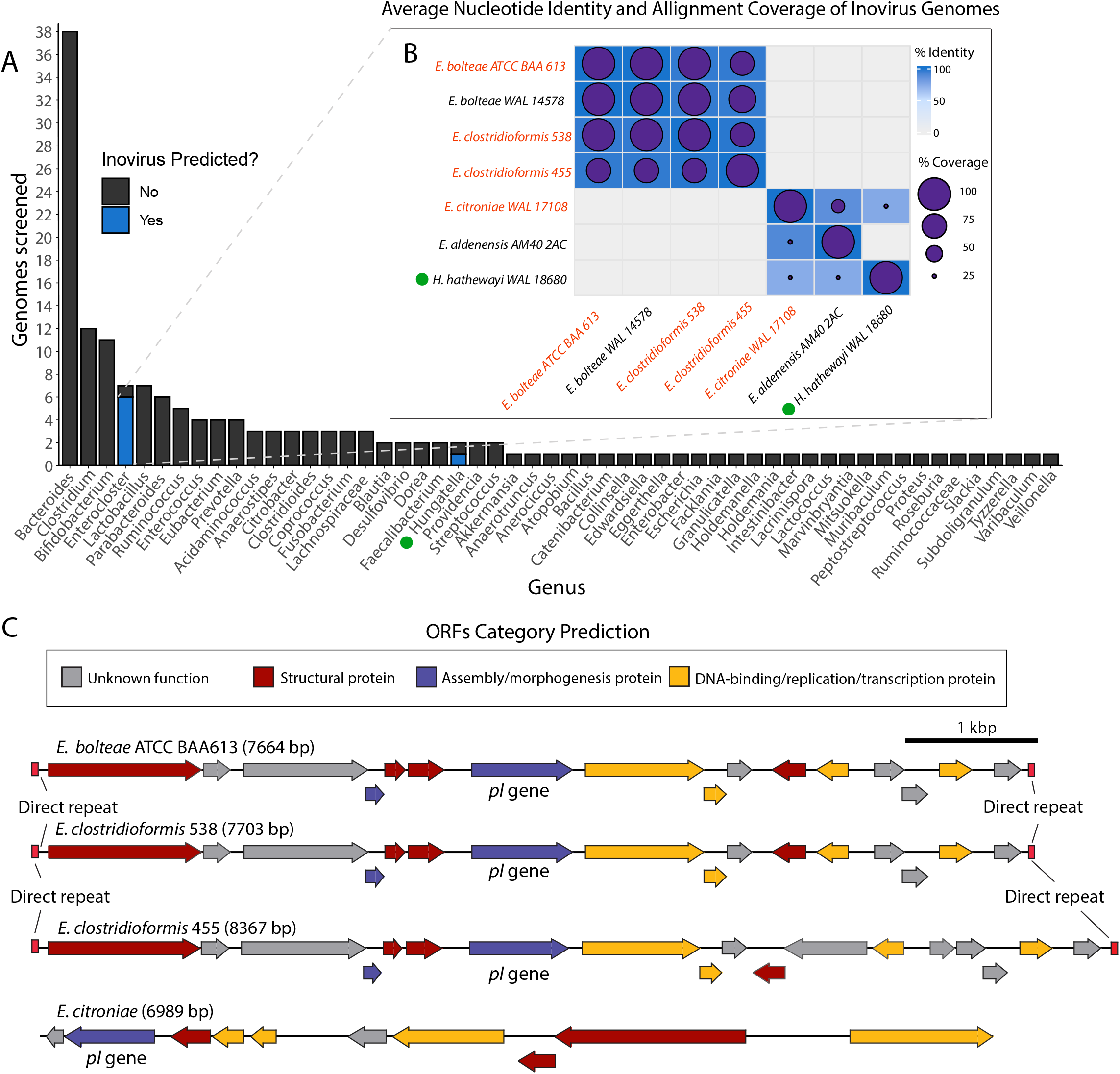
The genus *Enterocloster* harbors diverse inoviruses. **A)** Inovirus Detector [2] screen of 163 bacterial genomes [23] found putative inovirus prophages containing phage-related proteins in 6 strains of the *Enterocloster* genus and 1 strain of the *Hungatella* genus. Data was compiled based on genus; for a full list of bacteria screened, see Table S1. **B)** Average Nucleotide Identity (ANI) comparison of predicted inovirus genomes using the Pyani software [52] examine relatedness between genome sequences. Percentage sequence identity represented as the blue color gradient and sequence coverage by purple circles. Strains names colored in red were selected for downstream analyses and characterization. **C)** Linearized genomes from the four selected inovirus strains. ORFs were labeled based on general functions derived from HHpred predictions (see Table S5).

We performed an average nucleotide identity (ANI) analysis between the predicted inovirus genomes to examine their relatedness. Of the seven inovirus genomes we found, four of them were strongly conserved (>96% identity and ~73-100% alignment coverage) in *E. bolteae* and *E. clostridioformis* species and substrains, and the remaining three, found in *E. citroniae, E. aldenensis, and H. hathewayi*, shared few sequences (~10% coverage) with less conserved identity (73-90% identity) (Figure 1B). Interestingly, the few inovirus sequences shared between *E. citroniae, E. aldenensis, and H. hathewayi* inovirus genomes, as well as the lack of any similarity to *E. bolteae* and *E. clostridioformis* strains, suggests there are multiple types of inoviruses infecting *Enterocloster spp*. (Figure 1B).

When we compared the genomic location of ORFs within the *E. bolteae* and *E. clostridioformis* inoviruses, we saw a high degree of genome synteny between them (Figure 1C), further supporting that these two species share highly similar inoviruses. There was limited genome synteny between the *E. citroniae, H. hathewayi* and *E. aldenensis* inoviruses, which shared no synteny with the *E. bolteae* and *E. clostridioformis* inoviruses (results not shown). Since our data points to *Enterocloster* strains carrying more than one unrelated inovirus, we expanded our Inovirus detector screen to 24 publicly available *Enterocloster* genomes from NCBI (Table S3). Our screen found three other *E. clostridioformis* strains and one *E. bolteae* strain that had predicted inoviruses in their genomes with similar ANI to other strains of the same species (Table S2 and Figure S2). Interestingly, there were substrains of the original *Enterocloster* species we screened that had no predicted inoviruses in their genome, suggesting that inoviruses could be strain-specific, or that these isolates had not encountered the phage before.

As it was previously reported that inoviruses were produced by *Clostridium spp*. [2,9], we also screened 195 *Clostridium* genomes publicly available from NCBI with Inovirus detector (Table S3). We found two predicted inoviruses in gut commensal *Clostridium* species *C. butyricum* and *C. beijerinckii*, which were genetically distinct from each other and from the inoviruses we had previously identified (Table S2, Figure S2). We also wanted to determine if our predicted inoviruses shared any sequences with other previously sequenced inoviruses. Thus, we expanded our ANI analysis to compare our predicted *Enterocloster* inoviruses to 45 representative genomes from the *Inoviridae* phage family (Table S4). However, our analysis revealed no shared sequence identity between *Enterocloster* inoviruses and other representative genomes (Figure S3). Altogether, our results suggest the presence of a series of diverse, uncharacterized inoviruses in some gut-inhabiting species from the *Enterocloster* genus.

### Validation and characterization of *Enterocloster* inoviruses

Based on our bioinformatics results, we selected four predicted *Enterocloster* strains for further analysis and characterization of their respective inoviruses: *E. bolteae* ATCC BAA613 (*E. bolteae), C. clostridioforme* 2 1 49FAA (*E. clostridioformis* 455), *E. clostridioformis* WAL7855 (*E. clostridioformis* 538) and *E. citroniae WAL-17108* (*E. citroniae*). We used HHpred [28] (https://toolkit.tuebingen.mpg.de/tools/hhpred) to manually annotate gene functionalities in the inoviruses genomes. HHpred assigned annotations to 60-80% of ORFs of the *Enterocloster* inovirus genomes and identified structural, assembly, DNA binding, replication, and transcriptional regulation proteins (Figure 1C and Table S2). Many annotations were derived from commonly known inoviruses such as *P. aeruginosa* PF1 phage, *Xanthomonas* phage phiL, *Salmonella* phage IKe and *V. cholerae* CTXΦ phage (Table S5).

To confirm that the putative inovirus genomes detected in our selected *Enterocloster* strains were not ancestral remnants from previous phage infections, but were true actively secreted phages, we cultured these strains and performed PCR and negative transmission electron microscopy (TEM) to detect extracellular virions produced *in vitro*. Our PCR method used primers to amplify the inovirus *pI* gene from inovirus particles found in bacterial supernatants; intact bacterial cells were removed, and extracellular DNA was digested (Methods, Figure S4A). Virion DNA was detected in all supernatants tested (Figure S4B). For negative stain TEM, we serially concentrated cell-free spent media to enrich for inovirus particles and stained the resulting concentrate with 1% uranyl acetate (Methods). We observed filamentous structures in *E. bolteae* and *E. clostridioformis* 455 of around 500-700 nm in length and 6-10 nm in diameter, which are consistently sized with previously detected inoviruses (Figure 2).

**Figure 2.**
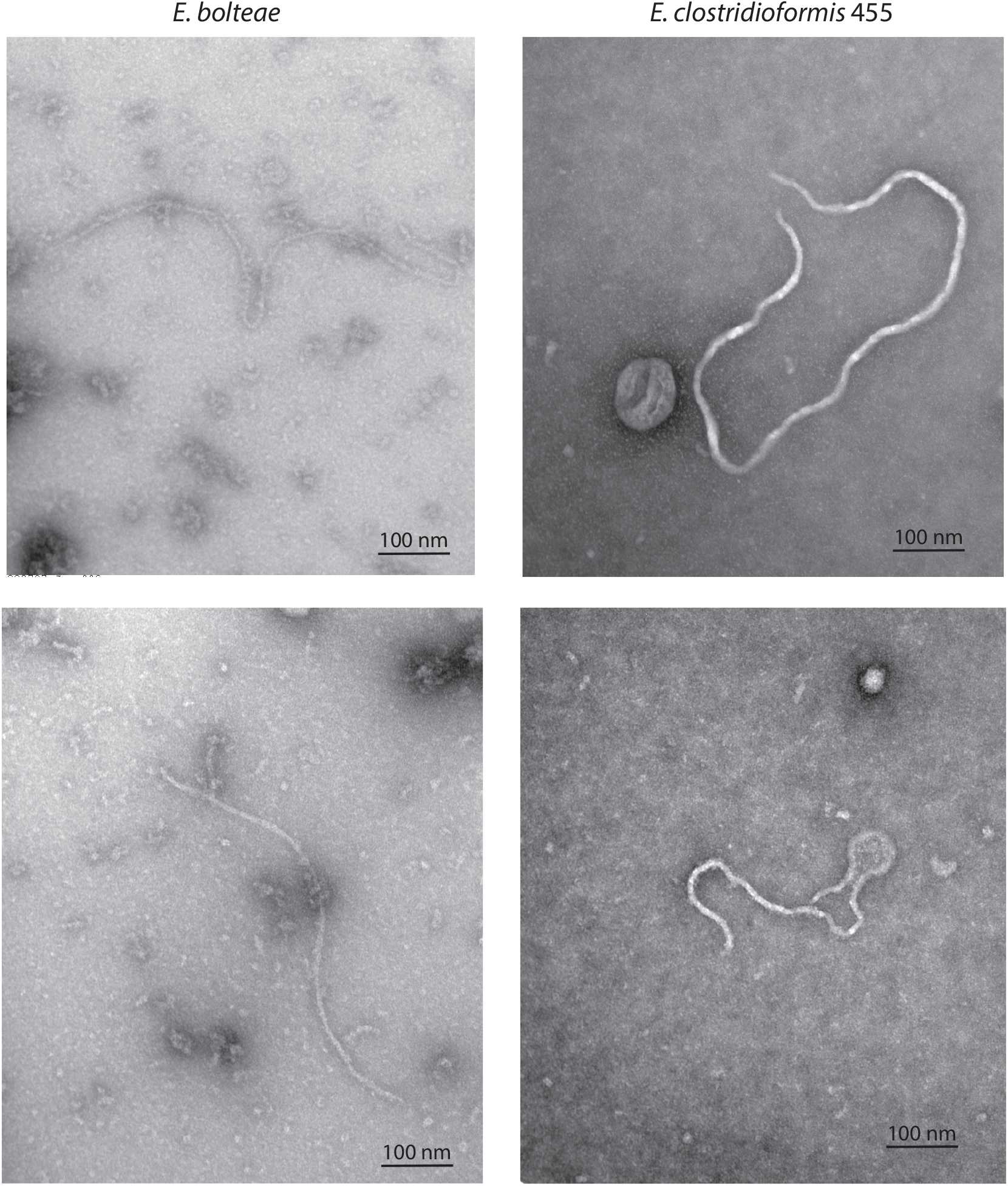
Inovirus virions are visible in *in vitro* cultures of *Enterocloster* strains. Negative stain transmission electron micrograph generated from concentrated and purified cell-free spent media supernatants of stable phase *Enterocloster* growth cultures. Specimens were stained with 1% uranyl acetate. Micrographs show filamentous morphologies with a diameter of ~6-8 nm and a length of ~600-800 nm.

### Inovirus secretion is affected by changes in environmental osmolality

Having confirmed inovirus production in *Enterocloster* strains, we investigated whether common environmental perturbations to the gut environment that impact the growth of *Enterocloster* may also affect inovirus secretion. *E. bolteae, E. clostridioforme* and *E*. *citroniae* are part of the Clostridia, a major class of mucosal colonizers of the microbiome (10-40% of total gut bacteria) that closely interact with the gut epithelium [29,30]. In previous work we had shown that changes in gut osmolality due to malabsorption strongly affect levels of Clostridia as well as phage membership in the gut [31]. We therefore assessed bacterial growth rate and inovirus secretion over a range of osmolality levels *in vitro*, that correspond to physiological conditions (~400-700 mOsm/kg) [31,32] (Methods, Figure 3A-D).

**Figure 3.**
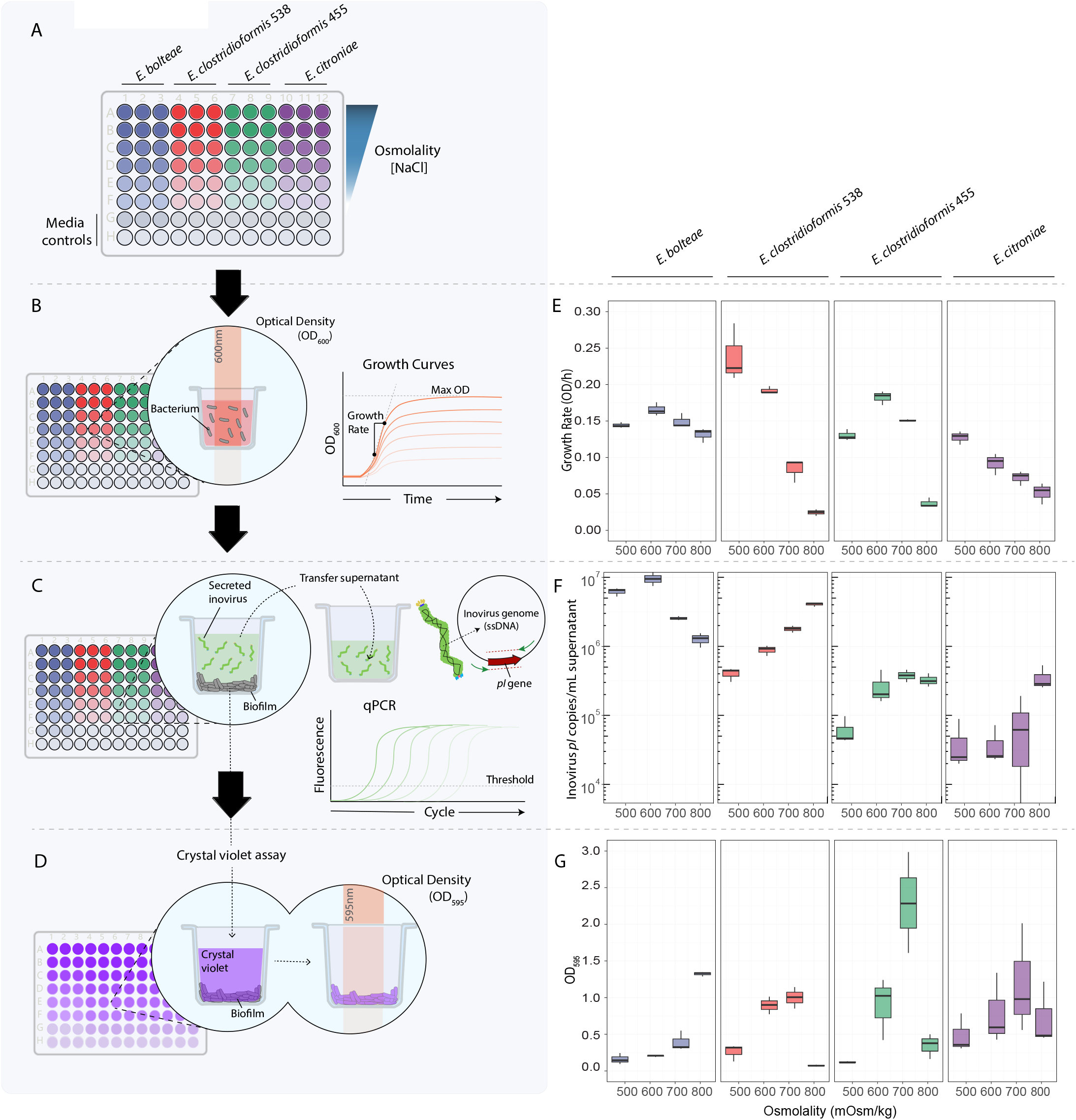
A multifaceted *in vitro* assay reveals species- and strain-specific inovirus production and biofilm formation in response to osmolality changes. **A)** A multifaceted *in vitro* assay to collect growth dynamics data and quantify copies of secreted inovirus and biofilm formation measurements. Strains were inoculated into a 96-well plate containing a range of different osmolalities. **B**, **E**) During a 63h growth period, OD_600_ measurements of each well were acquired every ten minutes using a plate reader. Growth curves were produced from the resulting OD_600_ data of each strain, and the bacterial growth rate (OD/h) was calculated by fitting the growth curves to a Gompertz curve. **C**, **F)** After incubation, the remaining cultures were transferred into a new plate, centrifuged, and the inovirus-containing supernatants were used to quantify secreted inovirus genome copy number by absolute qPCR. Inovirus genomes were detected using primers targeting the inovirus *pI* gene. **D**, **E)** A crystal violet stain was performed on the original culture plate, and OD_595_ measurements were taken to estimate biofilm formation in each well.

Under different levels of osmotic stress, we observed strain-specific changes in growth rate (Figure 3B,E) and overall maximum optical density (OD) (Figure S5), indicating varying and unique bacterial sensitivity to osmolality. We also saw diverse responses of inovirus secretion across the four *Enterocloster* strains tested in response to osmolality. Specifically, we saw a nearly 10-fold increase in inovirus secretion in *E. clostridioformis* 538 cultured at the highest osmolality (809 mOsm/kg) compared to baseline media (480 mOsm/kg), whereas increased osmolality did not promote inovirus secretion in *E. clostridioformis* 455 (Figure 3F). Furthermore, *E. citroniae* behaved similarly to *E. clostridioformis* 538, where increased osmolality promoted inovirus secretion (Figure 3F).

Since inovirus production is correlated with biofilm formation in other systems [16,19,33], we assessed biofilm formation using a crystal violet assay (Figure 3D). For example, Pf-4, the inovirus that infects *Pseudomonas aeruginosa*, is implicated in biofilm formation by promoting matrix crystallization [16,19]. Furthermore, osmotic stress has been reported to induce biofilm formation in many bacterial species [34], including those of the genus *Pseudomonas* [35,36]. However, we observed that biofilm formation was not broadly correlated with inovirus secretion for the strains tested (Figure 3F,G and Figure S6). For example, *E. bolteae* exhibited high biofilm formation at 809 mOsm/kg but reached both maximum inovirus secretion and growth rate at 622 mOsm/kg (Figure 3E,F,G). Together, our data suggest that the previous associations between biofilm formation and inovirus production [16] are not conserved for all inovirus-producing bacteria; instead, osmotic stress promotes inovirus secretion in a species- and strain-specific manner.

Given that inovirus secretion varied substantially between *Enterocloster* strains (Figure 3E), we assessed whether inovirus secretion patterns might be related to bacterial host physiology. To explore this, we investigated whether inovirus copies were correlated to maximum cellular growth rate in each osmotic condition (Methods). We found that inovirus secretion was positively correlated with growth rate in *E. bolteae*, negatively correlated in *E. clostridioformis*. 538 and *E. citroniae*, and not correlated in *E. clostridioformis 455* (Figure 4A). These results suggested that both in conditions where bacterial host growth is impaired or enhanced, inoviruses can still be produced at high levels in a strain-dependent manner.

**Figure 4.**
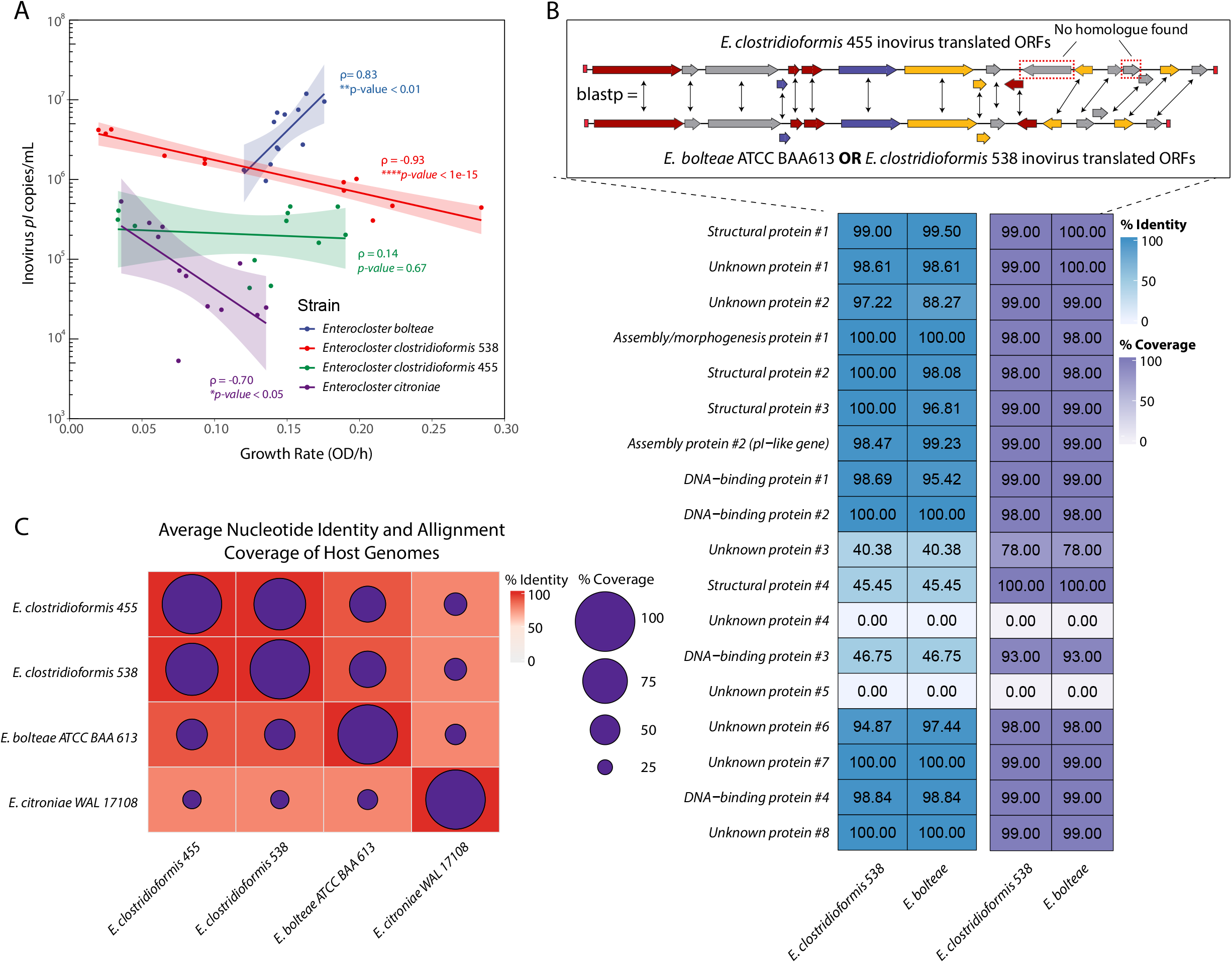
Related *Enterocloster* strains with highly similar inovirus prophages have unique relationships between inovirus secretion and host growth rate. **A)** Inovirus secretion and growth rate of *Enterocloster* bacteria were differentially correlated, depending on the *Enterocloster* species. Linear regression and 95% confidence intervals are shown for each strain. Correlation coefficients and *p-values* were obtained using Spearman correlation; * *p* < 0.05, ** *p* < 0.01, \*\*\*\**p* < 1×10^−5^ **B)** *top*: proteinprotein BLAST (blastp) comparisons of *C. bolteae* and *C. clost*. 538 translated ORFs against the *C. clost*. 455 translated ORFs; blastp top hit comparisons are depicted in the diagram with double-headed arrows. ORFs with no matches are highlighted by red, dashed boxes. *bottom*: resulting percentage identity and alignment coverage of blastp results. Row labels were assigned based on the annotation predicted in Figure 1C. Numbers in cells represent percentages. **C)** ANI comparison of *Enterocloster* host genomes. Percentage sequence identity represented as the red color gradient and sequence coverage by purple circles.

Interestingly, although our ANI data revealed that *E. bolteae, E. clostridioformis* 455 and *E. clostridioformis* 538 inoviruses were significantly similar (Figure 1B,C), we observed that these inoviruses were secreted at different levels in each host (Figure 4A). We compared the ORFs of their inoviruses to determine potential compositional differences that could explain the unique secretion patterns observed. Using protein BLAST (blastp), we compared the translated ORFs of *E. bolteae* and *E. clostridioformis* 538 against the translated ORFs of *E. clostridioformis* 455 (Figure 4B). Overall, we found that 13 out of 18 ORFs in *E. bolteae* and *E. clostridioformis* 538 closely matched an ORF in *E. clostridioformis* 455 (with ~98-100% identity and coverage) (Figure 4B). From the remaining 5 ORFs, 3 of them had partial matches to an ORF with low sequence identity (<47%) and 2 were unique to *E. clostridioformis* 455 (0% sequence identity). The 2 unique ORFs of *E. clostridioformis* 455 had unknown functions (Table S5). However, for the three proteins with partial matches, HHpred predicted a structural protein similar to the tail needle protein gp26, a DNA-binding omega regulatory protein and a protein with unknown function (Table S5). The differences between these proteins can be explored as potential explanations for the distinct secretion patterns observed in these *Enterocloster* strains. Surprisingly, when we compared the translated ORFs of *C. bolteae* against those of *E. clostridioformis* 538, we saw that all genes matched significantly (~90-100% identity and >98% coverage) (Figure S7), suggesting the possibility of other genes aside from those carried within inoviruses that may be involved in regulating their secretion.

Using ANI, we also tested the similarity between the *Enterocloster* host genomes (Figure 4C). *E. clostridioformis, E. clostridioformis* 455 and *E. clostridioformis* shared >91% sequence identity and ~50-90% alignment coverage, with while *C. citroniae* shared 78% sequence identity and 31-38% alignment coverage to the other strains (Figure 4C). The *E. bolteae* genome alignment coverage of ~51% against both *E. clostridioformis* 538 and *E. clostridioformis* 455 suggests that about half of the sequences in the *E. bolteae* genome are substantially similar to the genomes of these two strains (>91%), but the other half remain unique (Figure 4C). Our results suggest that the fewer host genome sequences shared by two inovirus hosts, the more different their inovirus secretion patterns may be. Altogether, there may be underlying host and inovirus genetic mechanisms involved in regulating inovirus secretion and thus specific host-inovirus pair should be investigated independently to assess secretion dynamics.

### Inovirus secretion is affected by osmotic stress *in vivo* in a species-specific manner

Osmolality is a major abiotic factor of the gut that impacts the composition of microbial communities in the microbiome [31]. Since we found osmolality had species-specific impact on inovirus production *in vitro*, we hypothesized that inovirus production may be similarly affected by osmotic changes in the gut. To test this hypothesis, we orally gavaged *E. bolteae* or *E. clostridioformis* 538 into germ-free mice (Methods). We selected these two strains as they contain an almost identical inovirus (Figure 1B,C and Figure S7), but differ in their optimal osmolality for *in vitro* inovirus secretion; peak *in vitro* inovirus secretion occur at different osmolality levels (~600 mOsm/kg in *E. bolteae* and ~800 mOsm/kg in *E. clostridioformis* 538, respectively) (Figure 3F). To increase gut osmolality, we added the common laxative polyethylene glycol (PEG) [31] into the drinking water of mice and, to achieve similar osmotic levels to the ones that induce peak inovirus production for these strains *in vitro*, we treated mice with two levels of PEG (10% and 15%, respectively) (Figure 5A, Methods). After six days of PEG treatment, we measured gut osmolality, bacterial abundance, and inovirus secretion in the cecal contents of mice (Figure 5B-E).

**Figure 5.**
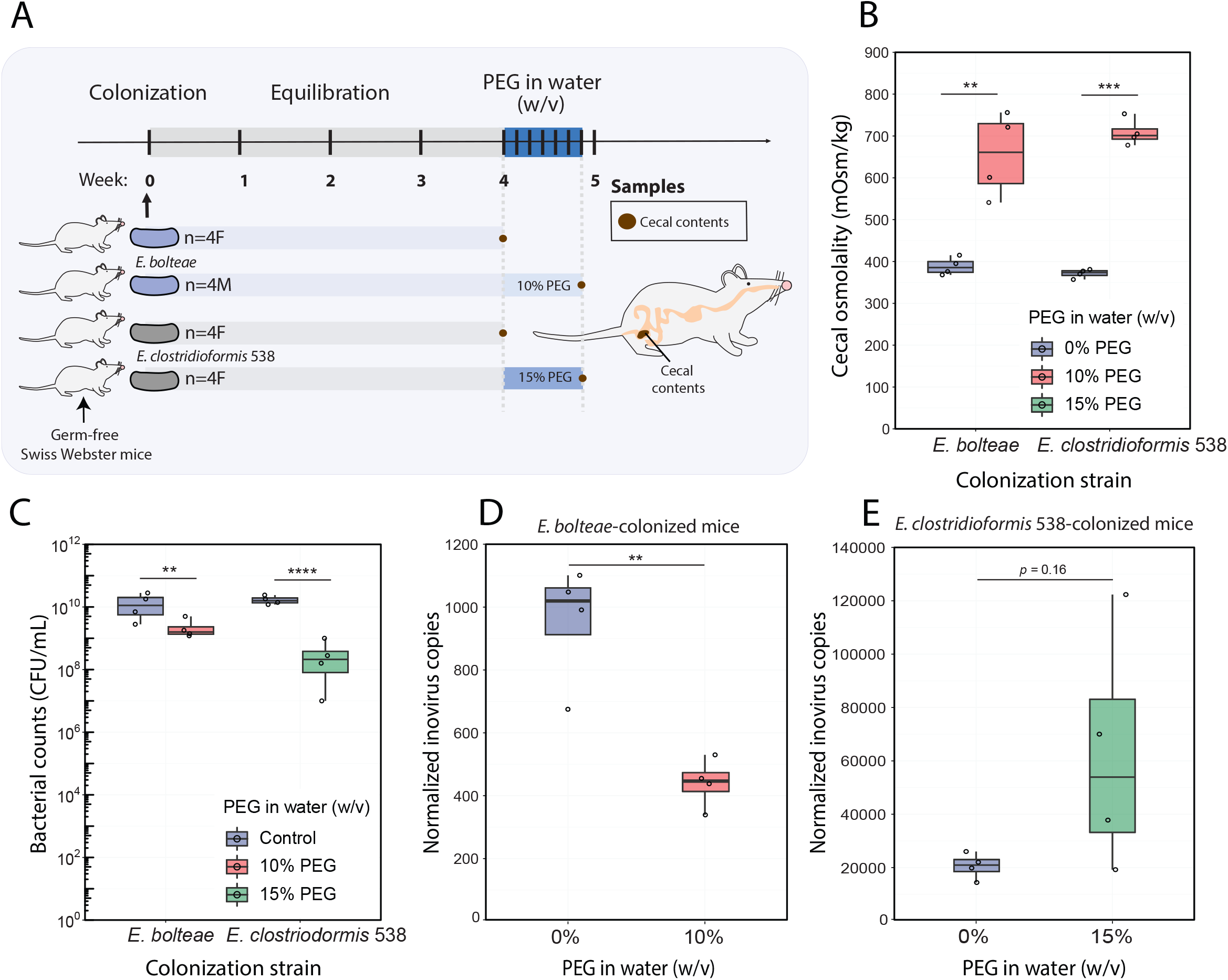
PEG treatment in *Enterocloster* mono-colonized mice impacts cecal osmolality, bacterial abundance and inovirus secretion in a species-dependent manner. **A)** experimental schematic: Germ-free Swiss Webster mice were gavaged with *E. bolteae* or *E. clostridioformis* 538, equilibrated for 4 weeks, and treated with 10% and 15% (w/v) PEG drinking water for 6 days, respectively. **B)** Cecal contents were collected from untreated and PEG-treated mice and centrifuged, and cecal osmolalities were measured from the resulting cecal supernatant. **C)** The effects of PEG treatment on bacterial density were measured by quantifying bacterial colony forming units (CFU/mL) from cecal contents. **D**, **E**) Relative qPCR quantification of inovirus production in cecal contents after PEG treatment. A primer pair was designed to quantify the conserved circularization region of the inovirus genome from *E. bolteae* and *E. clostridioformis* 538 using absolute qPCR (inovirus copies). Similarly, the unique integration sites of the inovirus genome in the *E. bolteae* and *E. clostridioformis* 538 genomes (gDNA copies). To account for changes in bacterial abundance after PEG treatment, inovirus copy numbers were normalized by the bacterial genomic copy number (inovirus copies/10^6^ gDNA copies). To test for significance, a Student’s t-test was performed between each experimental group and their control after accounting for group variances using Bartlett’s test; ** *p* < 0.01, \*\*\**p* < 0.001, \*\*\*\**p* < 1×10^−4^.

We found that cecal osmolality significantly increased in mice treated with 10% and 15% PEG. Median cecal osmolality was ~380 mOsm/kg for control mice that were not given PEG, and ~650 mOsm/kg and ~700 mOsm/kg for 10% and 15% PEG-treated mice, respectively (Figure 5B). Cecal bacterial counts (CFU/mL) for *E. bolteae* and *E. clostridioformis* 538 were roughly the same in untreated mice (~10^10^ CFU/mL), and both strains significantly decreased after PEG treatment (~10-fold for *E. bolteae*, and ~100-fold for *E. clostridioformis* 538) (Figure 5C). These results demonstrate that osmotic changes *in vivo* have a significant effect on the bacterial abundance of these *Enterocloster* strains. Furthermore, these results align with the decrease in growth rate and maximum OD we observed when these bacteria were cultured under osmotic stress *in vitro* (Figure 3E, S3).

To quantify inovirus produced in cecal contents, we used absolute qPCR. Given the linear ssDNA of secreted inoviruses circularize at specific regions, we designed primers that targeted and amplified only the circularized versions of the inovirus genome. This method allowed us to differentiate inovirus particles from integrated inoviruses prophages in the total extracted DNA from mouse cecal contents (Methods, Figure S8A). Additionally, to quantify bacterial genomic copy numbers using qPCR, we designed primers that amplified the unique integration sites of the inovirus genome in the *E. bolteae* and *E. clostridioformis* genomes (Methods, Figure S8B). Our qPCR data was consistent with the bacterial genomic copy numbers we quantified in the cecum via bacterial counts (~10^10^ copies/g of cecal content, Figure 5C, S9A). We also observed a significant drop in bacterial genomic copies after PEG treatment, consistent with cecal bacterial density data (Figure 5C, S9A). For *E. bolteae* mono-colonized mice, inovirus copy number started at 10^7^ copies/g of cecal contents—comparable to baseline inovirus expression *in vitro* (Figure 3F) — and decreased significantly (~10-fold) after PEG treatment compared to untreated controls (Figure S9B). Interestingly, we found that *E. clostridioformis* 538 produced more inoviruses in untreated mice than when grown to stationary phase in baseline media (Figure S9B, 3F), suggesting there might be specific factors influencing inovirus expression *in vivo*. To look at the secretion patterns of inoviruses in PEG-treated mice, we normalized the cecal inovirus copy number of each mouse by its respective bacterial genomic copy number (Figure 5D-E). Our normalized data determined that PEG treatment significantly decreased inovirus production in *E. bolteae* mice and increased inovirus production in *E. clostridioformis* 538 compared to untreated mice. This corroborated our *in vitro* observations of impaired inovirus secretion by *E. bolteae* and enhanced secretion by *E. clostridioformis* 538 at high osmolalities (Figure 3E).

## Discussion

Phages have been extensively shown to regulate the gut microbiota [37]; inoviruses in particular have been previously reported to be critical modulators of pathogen function [8,15,18–20]. Thus far, however, inoviruses have been understudied in the context of non-pathogenic gut bacteria. Previous metagenomic studies found that the Inoviridae family represents a small but significant fraction of the phages detected in the gut [2,21,38]. However, prior to our study, the provenance of these phages remained unknown. Our approach revealed that, within a representative list of gut commensals [23], inoviruses were mainly present in the *Enterocloster* genus (Figure 1A). Whether members from this genus are the primary gut commensals capable of secreting inoviruses remains to be determined. However, the lack of filamentous phages in the microbiota genera we surveyed raises questions about why these inoviruses are rare in gut commensals. A potential explanation is that the cost of inovirus reproduction exceeds the potential benefits conferred by the phage. For example, inovirus-producing *P. aeruginosa* grows slower than their non-infected counterparts, emphasizing a metabolic cost linked to inovirus infections [16,19]. Furthermore, Ff inoviruses infecting the Enterobacteriaceae were shown to impair host protein and RNA expression [39] and caused envelope stress [40,41], indicating additional negative impacts resulting from these phages. Bacteria of the gut microbiota face constant competition for nutrients and other resources: the excess energetic cost or adverse physiological effects resulting from inovirus hosting could select against bacteria harboring them.

Our analysis revealed that there are *Enterocloster* strains that do not carry inovirus prophages, indicating that inoviruses are not completely penetrant in the *Enterocloster* genus (Table S3). Interestingly, inoviruses appeared to be specific to gut-associated *Enterocloster* species, suggesting that the gut environment may be important for the retention and proliferation of these particular phages. The presence of inoviruses in a considerable fraction of *Enterocloster* gut bacteria evokes the question of what benefits, if any, are conferred by inoviruses to promote their conservation in these gut commensals. Previous studies have revealed that secretion of inovirus Pf-4 promoted phenotypes associated with tempered inflammatory responses in the lung mucosa helping *P. aeruginosa* evade immune detection [19]. *Enterocloster* commensals have been linked to immune tolerance in the gut [42], but the mechanisms behind this are still being uncovered. Since the intestinal lumen is a site of active interactions with the microbiota and the host immune system [43], it will be important to further investigate a possible role for inovirus secretion in immune tolerance of gut bacteria.

Interestingly, while all the *Enterocloster* strains we characterized were isolated from healthy human donors, annotations from the *E. bolteae, E. clostridioformis* and *E. citroniae* inovirus genomes returned hits to the zonulin occludens toxin (Zot) from the CTXΦ inovirus in *V. cholerae* (Figure 1B, Table S5). Beyond facilitating CTXΦ virion secretion, Zot functions as a mild toxin that affects epithelial tight junctions and increases gut permeability [18]. Previously, links between the enrichment of *Enterocloster* species and gastrointestinal diseases have been observed. While any potential toxicity of Zot-like proteins in *Enterocloster* inoviruses must be confirmed experimentally, these proteins could be tied to disorders associated with *Enterocloster* enrichment, and it will be important to include inviruses in the lens of future studies exploring these connections. For example, the correlation found by previous studies between enrichment of *Enterocloster* species, autism spectrum disorder and gastrointestinal diseases [22,44,45] should be further explored in the context of inovirus presence.

Beyond the genomic characterization of these novel inoviruses, in this study we tested how osmotic perturbation affects inovirus production. Previous studies reported that *E. coli* is more sensitive to heat shock, osmotic shock, and freeze-thaw cycles when infected with M13 inovirus [46]. We observed species-specific patterns of inovirus secretion linked to osmotic changes *in vitro* within members of the *Enterocloster* genus (Figure 3). Despite previous observations showing that inovirus secretion was related to decreased growth rate [16,19,46,47], we found that changes in growth rate were not consistently correlated with inovirus secretion in the *Enterocloster* strains tested (Figure 4A). Given that phage transporters are inserted in the envelope of the bacterial host, the combination of osmotic shock and of phage secretion may affect bacterial viability in a species-dependent manner. Altogether, exploring the effects of physical factors in inovirus secretion, especially in a dynamic and heterogeneous environment like the gut [48,49], will be critical in understanding the function of *Enterocloster* inoviruses in the microbiota.

Despite previous observations showing that secretion of Pf-4 inoviruses in *P. aeruginosa* facilitate biofilm formation [16,20], we found no significant correlation between biofilm formation and inovirus secretion in any bacteria and osmotic conditions tested (Figure 3, S6). Secretion of Pf-4 inoviruses facilitates biofilm formation by interacting with the lipids, proteins, and lipopolysaccharides of the Gram-negative bacterial outer membrane [50]. The Gram-positive nature of *Enterocloster spp*., lacking an outer membrane to interact with inoviruses, could explain why we saw no relationship between biofilms and inovirus secretion. However, because only two cases of Gram-positive organisms producing inoviruses have been reported [9–11], host-inovirus interactions remain poorly understood in Gram-positive organisms. Exploring structural or mechanical interactions between inoviruses and the Gram-positive cell walls of *Enterocloster spp*. could reveal important functionalities for these phages.

Many questions remain about the origin of *Enterocloster* inoviruses and their potential function, and their niche in the context of the microbiota. The increasing availability of gut viral genomic and metagenomic datasets can help uncover how widespread *Enterocloster* inoviruses are in the microbiota and in which conditions they are most represented (*e.g*., during dysbiotic and pathological conditions). Furthermore, the presence of specific inoviruses in gut samples can be quantified using the qPCR methods outlined in this study, allowing for a more quantitative and sensitive interrogation of these phages in the microbiota. Overall, understanding what role inoviruses may play in commensal bacteria may enable us to understand how major microbiome members adapt and persist in the gut microbiome.

## Conclusion

Our characterization of endogenous gut symbionts harboring inoviruses counters the existing assumption that inoviruses in humans are reserved for Gram-negative pathogenic bacteria. Our research reveals that the host range of these filamentous phages may be more extensive than currently predicted. These phages may occupy other non-pathogenic niches in the gut microbiome and may have implications for the opportunistic pathogenicity of their hosts. Determining how common, or in which cases, *Enterocloster* inoviruses are present in the gut will help us grasp how important these phages are in the microbiota. Furthermore, the functional and structural characterization of *Enterocloster* inoviruses will be key to deriving new hypotheses about the niche these filamentous phages fill in the gut. Overall, understanding the effect of inoviruses in *Enterocloster spp*. will expand our understanding of *Enterocloster* biology and its roles in the microbiota.

## Materials and methods

### Bacterial cultures and media preparation

All bacteria were grown anaerobically (85% N_2_, 10%CO_2_, 5% H_2_) at 37°C in a modified Brain Heart Infusion (BHIS-YE) medium. This medium contains 37g/L Brain Heart Infusion (BD Biosciences) supplemented with 5 mg/L of hemin, 2 mg/L of vitamin K1 and 5g/L of yeast extract. Solid agar plates were prepared by combining Brain Heart Infusion, yeast extract, and 15 g/L agar and then autoclaving for 30 minutes at 121°C. Once cooled, the agar was supplemented with hemin (5 μg/mL) (MilliporeSigma) and vitamin K1 (1 μg/mL) (Alfa Aesar) and plates were poured. For liquid media only (BHISG-YE), dextrose was added to a final concentration of 100 mM to promote biofilm formation and MgSO_4_ and CaCl_2_ was added to a final concentration of 1 mM to promote phage adhesion [51]. All components were dissolved in MilliQ water and filter sterilized. Overnight cultures used in all experiments were grown statically in 6-well plates with 3 mL of BHISG-YE media.

Columbia blood agar plates were used to plate *in vivo* samples. This medium consists of 35g/L of Columbia Broth Powder (BD Difco™) dissolved in water and autoclaved 30 minutes at 121°C. Once cool, and before pouring plates, defibrinated sheep blood (Hemostat Laboratories) was added to the media at a final volume of 5% (v/v), and supplemented with hemin (5 μg/mL) (MilliporeSigma) and vitamin K1 (1 μg/mL) (Alfa Aesar).

### *In silico* detection of inovirus genomes

Inovirus Detector by Simon Roux [2] was used to predict inoviruses in the genomes of bacteria. Software and documentation is published by Simon Roux on Github (https://github.com/simroux/Inovirus). The NCBI datasets command line tool v13.21.0 was used to download genomes using the Assembly accession ID for the different strains. Putative inovirus genomes were extracted from their respective host genome file using the genome coordinates given by the inovirus detector output. All genomes extracted can be found in FASTA format in the supplementary information section.

### ANI comparisons

Average Nucleotide Identity (ANI) comparisons (sequence identity and alignment coverage) of genomes were performed using Pyani v0.2.11 [52] using default settings except for the method (-m) where ANIb was used as an argument. Percentage identity and alignment coverage output files were parsed using packages tidiverse v1.3.2 and readxl v1.4.0 and plotted using ComplexHeatmap v2.10.0

### Inovirus genome annotations

Homology-based searches (BLAST and PFAM) were used for *Enterocloster* inovirus ORFs to predict functional annotations but had limited success. HHpred [28] (https://toolkit.tuebingen.mpg.de/tools/hhpred), a software better suited for remote homolog prediction, was implemented to manually annotate gene functionalities of selected inovirus genomes. Top HHpred predictions were used to annotate the ORFs.

### PCR detection of inoviruses *in vitro*

A single colony from each bacterial species was inoculated in 3 mL of BHISG-YE in separate wells of a 6-well plate. The next day, an additional 3 mL of BHISG-YE was added. Every other day, 3 mL of bacterial culture was removed from each well and replaced with 3 mL of fresh BHISG-YE. On the ninth day, 1 mL of culture was harvested from each well and centrifuged spun down for 10 minutes at 5,000g to pellet bacterial cells. The supernatants were then passed through a 0.22-micron filter. In a PCR tube, 14 μL of filtered supernatant was combined with 4 μL of RDD buffer (Qiagen, RNase-Free DNase Set) and 2 μL of DNase I and dissolved as per the manufacturer’s instructions. Supernatants were then incubated at 37°C for 1.5 hours and then heated at 85°C for 15 minutes to denature DNase and open inovirus capsids.

Immediately after DNase denaturation step, PCR was performed on 1 μL of supernatant using primer sets targeting the pI gene in *E. citroniae* (F: 5’-TCGGTTCATCACTGCGTAAG-3’, R: 5’-GGTAGATGGCGAGGTTGTTG-3’) and in the conserved pI gene of *E. bolteae, E. clostridioformis 455* and *E. clostridioformis* 538 (F: 5’-CCAGGCGTATCACAAAGACA-3’, R: 5’-CAGGAGCAGGGAATCAATGT-3’). To ensure that amplification of the *pI* gene was not derived from bacterial chromosomal DNA, amplification of the V7-V8 region of the 16S rRNA gene using universal primers 1237F (5’-GGGCTACACACGYGCWAC-3’) and 1391R (5’-GACGGGCGGTGTGTRCA-3’) were also used on the supernatant samples. PCR reactions were performed using DreamTaq Green PCR Master Mix (ThermoFisher Scientific) and final primer concentrations of 200 nM. Thermal cycling was performed for 35 cycles with an annealing temperature of 52°C. PCR products were run on a 1.5% ethidium bromide-stained agarose gel and product band mean density was quantified using ImageJ. Background mean density was subtracted from wells containing no template controls for each primer set at the expected location of the DNA product band. Mean densities are plotted in Figure S4B.

### Negative stain transmission electron microscopy

#### Sample preparation

Single colonies were inoculated into 3 mL of BHISG+YE and incubated overnight anaerobically at 37°C. Overnight cultures were then subcultured 1:100 into 40mL of ~620mOsm/kg (*C. bolteae*) or ~820 mOsm/kg (*E. clostridioformis* 455) BHISG+YE media and incubated anaerobically at 37°C for 48 h. Cultures were then spun down at 4,000xg for 15 minutes to pellet cells, and 15 mL of supernatants were transferred into a 30 kDa Amicon® Ultra-15 Centrifugal Filter Units (Sigma). The residue from the filtration device was then transferred to a 1.7 mL microfuge tube and spun at 12,000xg for 5 minutes to pellet any remaining cells. The supernatants were transferred to 1.7 mL microfuge tubes and TE buffer was added to reach 1 mL. 200 uL of a 20% PEG 6000 and 2.5 M NaCl solution were added to each supernatant, mixed, and incubated for 15 minutes at room temperature. The supernatants were then spun at 12,000xg for 10 min to pellet the inoviruses, the excess media was removed, and then spun again at 12,000xg for 5 min. The excess media was removed, and the pellets were resuspended in 500 uL of TE buffer. Resuspended pellets were transferred into a 10 kDA Amicon Ultra-0.5 Centrifugal Filter Unit, spun at 14,000g for 15 minutes, and the concentrate was collected and stored at 4°C until use.

#### Negative stain TEM

3 μl of each concentrated supernatant were put onto a glow discharged grid and incubated for 3 min. After removing the drop of concentrated supernatant with filter paper, 3 μl of 1% uranyl acetate were placed on to the grid as the negative stain for 30 seconds. Negatively stained samples were observed using an FEI Tecnai Spirit transmission electron microscope operating at 120 kV.

### Growth curves coupled with biofilm and PCR inovirus detection assays

For experiments in Figure 3 and Figure S4, BHISG-YE media osmolality was measured on the Advanced Instruments Model 3320 Osmometer and adjusted using a 1M NaCl BHISG+YE media to 622 mOsm/kg, 723 mOsm/kg and 809 mOsm/kg. The baseline osmolality of BHISG-YE is 480 mOsm/kg. Triplicate wells containing 200 μL of each media condition were dispensed in a 96-well plate and reduced in anaerobic conditions for 12 hours.

Overnight cultures of each bacterium were refreshed by adding 1 mL of BHISG-YE and incubated at 37°C for 6 hours prior to seeding. OD_600_ of the refreshed culture was measured on a microplate reader (BioTek Synergy H1) contained in an anaerobic chamber and diluted with BHISG-YE to a final OD_600_ of 1 before seeding 2 μL of culture in triplicate wells of each media condition in a 96-well plate. Well plates were sealed using a plastic seal with holes poked with a sterile syringe and grown in an anaerobic plate reader for 63 hours with OD_600_ being measured every 10 minutes. A breathable seal was necessary, as all four bacteria appear to not grow without gas exchange (results not shown). Sterile blank wells of each media type were run in quadruplicate, averaged, and to subtracted from the OD_600_ readings of seeded wells. Growth rates were interpreted as OD/hr measurements by fitting a Gompertz curve to the exponential region through manual identification using an in-house MATLAB program.

After 63 hours of growth, cultures were gently aspirated and transferred to a separate 96-well plate that was subsequently centrifuged at 8000 RPM for 15 minutes. Meanwhile, a crystal violet assay (below) was performed to quantify the biofilm on the original 96-well culture plate.

After centrifugation, 18 μL of the supernatant was carefully removed from the top of the wells and placed in a 96-well PCR plate with wells containing 2 μL DNase I-RDD buffer mix. Supernatants were sealed and incubated at 37°C for 1.5 hours and then heated in a thermocycler to 85°C for 15 minutes in order to denature DNase and open inovirus capsids. After heat denaturing, samples were separated to individual well plates according to species. These plates were sealed with an aluminum seal and then frozen at −80°C. Separating samples by species guaranteed that all samples within a species were thawed for the same amount of time and not subjected to multiple freeze-thaw cycles as real-time qPCR was performed on each species across two separate days.

Real-time qPCR was performed using the PerfeCTa SYBR Green SuperMix (Quantabio) on the supernatant samples. For each sample, primer sets (above) targeting both pI and 16S genes were used, and standard curves were generated using known amounts of purified bacterial genomic DNA. All samples had Cq values for 16S genes below the no template controls, therefore are declared to be free of bacterial chromosomal DNA contamination.

### Biofilm quantification using crystal violet assays

Immediately after aspirating cultures for centrifugation and DNase treatment (above), wells were gently washed using 300 μL of PBS and placed upside down to dry for 20 minutes. 300 μL of 0.1% crystal violet solution was added to each well and incubated at room temperature for 10 minutes. Wells were then aspirated and washed three times with PBS and allowed to dry overnight. To dissolve the dried crystal violet, 300 μL of 33% glacial acetic acid was gently mixed into each well and OD_595_ was read in the plate reader.

### Protein-protein BLAST ORF comparisons

Putative inovirus ORFs were extracted from their respective host genome file using the ORFs coordinates given by the inovirus detector output. Nucleotide CDSs were translate to amino acids sequences using the EMBOSS transeq web page (https://www.ebi.ac.uk/Tools/st/emboss_transeq/) and stored as protein FASTA files. The BLAST command-line tool v2.10.1 was used to create local BLAST databases using the protein FASTA files. These local databases served as a reference to compare the protein ORFs from the other strains using protein-blast (blastp). Query sequence identity and alignment coverage were extracted from the top hit of each ORF. All CDS files can be found in FASTA format in the supplementary information section.

### *In vivo* experiments

All animal experiments were performed in accordance with the University of British Columbia Animal Care Committee. In this study, Swiss-Webster germ-free (GF) mice were maintained in gnotobiotic isolators until the experimental age of 4-5 weeks. Once experimental age was reached, littermates of the same sex were aseptically transferred into four gnotobiotic isocages in groups of four. Mice were then colonized by oral gavage with 200 μL of *Enterocloster* bacterial cultures in stationary phase. Two cages of female mice (4 per cage) were colonized with *E. clostridioformis 538*, and two cages, one with female mice and the other with male mice (4 per cage), were colonized with *E. bolteae*. Mice were fed an autoclaved standard diet (Purina LabDiet 5K67). After four weeks of equilibration, control cages (female mice) of *E. bolteae* and *E. clostridioformis 538* colonized mice were sacrificed, and their cecal contents were collected. For the remaining cages, 10% and 15% of PEG 3350 (Miralax) was added to the drinking water of the *E. bolteae* and *E. clostridioformis* 538 colonized mice, respectively. Mice were given PEG water for six days before sacrificing and collecting cecal contents.

*To measure cecal osmolality*, cecal contents were centrifuged at 16,000x*g* for 20 mins at 4°C after collection. The resulting cecal supernatant was used to measure osmolality as above.

*To quantify bacterial abundance*, cecal bacterial colony forming units (CFU) were quantified by sampling with 1uL loops, followed by 10-fold serial dilutions and spot plating 5 μL, in duplicates, on both BHISG+YE (for counting) and Columbia blood plates (to assess for contamination). Plates were incubated anaerobically for 1-2 days at 37°C until visible colonies were observed.

### Absolute quantification of inoviruses from cecal samples

Cecal DNA was extracted from ~30-200 mg samples using the DNeasy PowerSoil Pro Kit (Qiagen). The concentration of extracted DNA was measured with a NanoDrop Lite (Thermo) spectrophotometer, and the DNA from all extracted samples was normalized to 4 ⊓g/μL.

For absolute inovirus and bacterial genome copy number quantification, real-time quantitative PCR (qPCR) was performed with 25 μL reaction volume using PerfeCTa SYBR Green FastMix (Quantabio); 300 nM final primer concentration per well and 20 ng of extracted DNA per well were used for all qPCR reactions. Thermal cycling was performed in a C1000 Touch Thermal Cycler (Biorad) with the following conditions: initial denaturation of 95°C for 2 mins, 35 cycles of 95°C for 20 sec, 58°C for 30 sec, and 72°C for 20 sec. For standard curves, threshold cycle (Ct) values demonstrated a linear dependence (R^2^ = 0.99) to the standard concentration value, and PCR efficiency ranged between 90-110%.

*Absolute Inovirus genome quantification* was determined using a standard curve method and a primer set targeting the conserved circularization region of the *E. bolteae* and *E. clostridioforme* 538 single-stranded DNA inovirus genomes (F: 5’-TTGCTGACGCCCTCTCTGAC-3’, R: 5’-CACCCCCTAACAAAAGTGTTAAAAG3’). The standard curve was created from PCR product generated using the inovirus primer set (above) and a Phusion® High-Fidelity DNA Polymerase (NEB) using extracted DNA from each strain as a template. PCR product was purified using the QIAquick PCR Purification Kit (Qiagen), quantified via nanodrop, and used as stock for a 10-fold serial dilution standard curve. Based on the generated calibration curves, the resulting sample Ct values were converted to inovirus copies/g of cecal content.

*Absolute bacterial genome quantification* was determined using a standard curve method and a primer set that spanned the bacterial genome and the inserted inovirus genome, unique to each strain (*E. bolteae*, F: 5’-ATTGCTGACGCCCTCTCTGAC-3’, R: 5’-AAGAACAAGGAACCTCACCCC-3’; *C. clostridioforme 538*: F: 5’-ATTGCTGACGCCCTCTCTGAC-3’, R: 5’-CGCAGGGATTACAAAGACTAACCC-3’). Standard curves were generated using known amounts of purified bacterial genomic DNA (gDNA) extracted from stable phase cultures using the DNeasy PowerSoil Pro Kit (Qiagen). Purified gDNA was quantified via nanodrop and used as stock for a 10-fold serial dilution standard curve. Based on the generated calibration curves, the resulting sample Ct values were converted to gDNA copies/g of cecal content.

*Normalization of inovirus genome copies* was calculated by dividing the inovirus genome copy number by the gDNA copy number for the same sample and multiplying by 1,000,000.

## Acknowledgements

The authors acknowledge that the land we performed this research on is the traditional, ancestral, and unceded territory of the xwməθkwəýəm (Musqueam) Nation. We encourage others to learn more about the native lands in which they live and work at https://native-land.ca/. This work also received support from the Bioimaging facility (RRID: SCR_021304) at the University of British Columbia.

This research was supported in part through computational resources and services provided by Advanced Research Computing at the University of British Columbia. We acknowledge Dr. Katharine Ng for useful discussion, help with experimental design and providing feedback for this manuscript, Dr. Samuel Collins and Negin Rahanjam for support in animal husbandry, and Giselle McCallum for critically reading this manuscript and providing feedback. We also thank Amy Chan for her support and feedback in negative transmission electron microscopy protocols. The authors acknowledge support from the Natural Sciences and Engineering Research Council of Canada Discovery Grant (to C.T.) and from the 4 Year Fellowship scholarship from UBC (Juan C. Burckhardt).

## Supplementary figures and tables legends

**Figure S1. Pie chart of the phyla represented by the 163 gut bacteria screened with Inovirus detector.** See Table S1 for the complete list of bacteria.

**Figure S2. Expanded ANI analysis of inoviruses.** Average nucleotide Identity (ANI) comparison of putative inovirus genomes (Table S2). This comparison includes identified inovirus genomes found after screening the *Clostridium* (green) and *Enterocloster* (purple) genomes in Table S3, as well are the original inoviruses we identified (red) (Figure 1B). To further screen the *Clostridium* and *Enterocloster* inovirus genomes against a larger reference database, we performed nucleotide BLAST searches of the inovirus genomes and tBLASTx searches of their ORFs (results not shown). Other than to themselves, the BLAST matches were not significant, indicating a high specificity of these inoviruses for their strains.

**Figure S3. Putative inoviruses share no sequence identity to other reference inoviruses.** ANI comparison of 45 inovirus genomes (Table S3) downloaded from NCBI and predicted inoviruses found in this study (highlighted in red).

**Figure S4. Inoviruses are secreted *in vitro*. A)** A schematic demonstrating how inoviruses are detected *in vitro*. First, nine-day-old cultures were separated to yield cell-free spent media supernatants that were filter-sterilized and then treated with DNase to eliminate contaminating bacterial DNA. The DNase-treated supernatant was then used as template DNA in PCR reactions using primers targeting the pI gene and 16S rRNA gene to guarantee that there was no 16S contamination. **B)** Bands resolved by gel electrophoresis from amplified DNase treated supernatants (cream), and positive control purified genomic DNA (blue) were measured using ImageJ with background subtracted in corresponding water-control lanes for *E. clostridioformis* 455, *E. clostridioformis* 538, *E. citroniae*, and *E. bolteae*. Selective detection of DNase-resistant *pI* genes suggests inoviruses are secreted in vitro. Genomic DNA control is not representative of physiological copies of *pI* and 16S because PCR was performed using purified DNA with 10 ng of template, which likely saturated the reaction.

**Figure S5. Overall maximum OD_600_ of *Enterocloster spp*. change in a strain-specific manner in response to osmotic stress.** Maximum OD_600_ of selected *Enterocloster* strains was calculated from the growth curves obtained as explained in Figure 3A-B. Under different osmotic stresses, we observed varying bacterial sensitivity to osmolality at both the strain and species levels. Notably, we saw a decrease in growth rate (Figure 3E) and overall max OD when osmolality was increased above the baseline media osmolality of 480 mOsm/kg in *E. clostridioformis* 538 and *E. citroniae*. Unlike its counterpart strain, we noticed that the slower-growing *E. clostridioformis* 455 exhibited tolerance to increased osmolality up to 722 mOsm/kg in the form of stable growth rates, despite decreasing overall max OD (Figure 3E). Uniquely, *E. bolteae* exhibited faster growth rates only at 622 mOsm/kg (Figure 3E), suggesting an optimal osmolality, despite declining overall max OD (Figure S3). Nonetheless, *E. bolteae* also showed the overall highest tolerance to osmolality from all tested strains, as both its growth rate and max OD were the ones that changed the least compared to baseline. These results highlight the unique relationships that bacteria, even at the species or substrain level, have with osmolality, which is consistent with previous findings in other gut bacteria [31].

**Figure S6. Inovirus secretion is not correlated with biofilm formation in *Enterocloster* strains.** Linear regression and 95% confidence intervals are shown for each strain. Correlation coefficients and *p-values* obtained using Spearman correlation.

**Figure S7. *E. bolteae* and *E. clostridioformis* 538 share significant amino acid contents of ORFs.** *Top*: protein blast (blastp) comparisons of *E. bolteae* translated ORFs against *E. clostridioformis* 538 translated ORFs; blastp top hit comparisons are depicted in the diagram with double-headed arrows. *Bottom*: resulting percentage identity and alignment coverage of blastp results. Row labels were assigned based on the annotations predicted in Figure 1C. Numbers in cells represent percentages.

**Figure S8. Validation of *in vivo* qPCR primers. A)** *top*: diagram depicting the circularization region targeted by primers to detect inovirus genomes, *bottom*: total DNA was extracted from stable phase *Enterocloster* cultures grown at the highlighted media osmolality. Extracted DNA was used as a template in PCR reactions containing the inovirus-specific primers. The resulting PCR product was resolved in a SYBR-stained 1% agarose gel. A 100 bp ladder was run in parallel to determine product size. **B)** *top*: diagram depicting the integration region amplified by primers within the inovirus integrated genome and the host genome to quantify gDNA copies. *bottom*: total DNA was extracted from stable phase *Enterocloster* cultures grown at base osmolality. Extracted DNA was used as a template in PCR reactions containing the gDNA primers. The resulting PCR product was resolved in a SYBR-stained 1% agarose gel. A 100 bp ladder was run in parallel to determine product size.

**Figure S9. Bacterial and inovirus genome copies decrease *in vivo* after PEG treatment. A)** Bacterial gDNA copies and **B)** Inovirus genomes copies quantified from cecal contents using absolute qPCR. To test for significance, a Student’s t-test was performed between each experimental group and their control after accounting for group variances using Bartlett’s test; ** *p* < 0.01, \*\**p* < 0.001.

**Table S1. List of genomes modified from Han *et al*.** [23] **that were screened with Inovirus detector.** Bacteria whose taxonomy has changed since the *Han et al*. paper are shown in red; those pending approval to change their taxonomy are shown in blue.

**Table S2. Strains with putative inoviruses**

**Table S3. List of *Enterocloster* and *Clostridium* genomes downloaded from NCBI.**

**Table S4. List of representative inovirus genomes downloaded from NCBI.**

**Table S5. HHpred annotations for selected *Enterocloster* inoviruses.**

